# High carnivore population density highlights the conservation value of industrialised sites

**DOI:** 10.1101/260729

**Authors:** Daan J. E. Loock, Samual T. Williams, Kevin W. Emslie, Wayne S. Matthews, Lourens H. Swanepoel

**Author notes:** These authors contributed equally to this work, and are listed in alphabetical order (Daan J. E. Loock, Samual T. Williams).

## Abstract

As the environment becomes increasingly altered by human development, the importance of understanding the ways in which wildlife interact with modified landscapes is becoming clear. Areas such as industrial sites are sometimes presumed to have little conservation value, but many of these sites have areas of less disturbed habitats around their core infrastructure, which could provide ideal conditions to support some species, such as mesocarnivores. We conducted the first assessments of the density of serval (*Leptailurus serval*) at the Secunda Synfuels Operations plant, South Africa. We ran three camera trap surveys to estimate serval density using a spatially explicit capture recapture framework. Servals occurred at densities of 76.20-101.21 animals per 100 km^2^, which are the highest recorded densities for this species, presumably due to high abundance of prey and the absence of persecution and/or competitor species. Our findings highlight the significant conservation potential of industrialised sites, and we suggest that such sites could help contribute towards meeting conservation goals.

## 1 Introduction

Over the last centuries, there have been rapid and intense environmental changes caused by increasing human numbers, technological advances and industrialisation (United Nations Environment Programme 2012). Human alterations on the environments have resulted in a decline in biodiversity, and are elevating extinction rates of species at a global scale (Chapin et al. 2000). Currently more than 75% of the terrestrial surface is impacted by humans (Ellis et al. 2010; Ellis et al. 2013). These human activities are affecting biodiversity and ecosystems on various scales as well as modifying existing habitats, creating unique urban environments and novel ecosystems (Hobbs et al. 2006; Williams et al. 2009; Barbosa et al. 2010). In many cases, biodiversity can be positively related to human population at a regional scale due, for instance, to an enhanced spatial heterogeneity between rural and urban environments, and the introduction of exotic species (McKinney 2002; Sax and Gaines 2003). The influence of these modifications depends on both the scale and the organisms involved (Barbosa et al. 2010).

Even within the most densely populated and intensively used areas, including urban landscapes, humans rarely utilise all land, and tend to retain significant green or unused areas. These “green spaces” hold ecological potential, and can reduce biodiversity loss by managing habitats to support endangered species (Jackson et al. 2014), although, further research is necessary to understand the impacts of these processes (Northrup and Wittemyer 2013) transformed landscapes lead to unpredicted changes in species communities, posing new challenges to conservation and resource management (Lindenmayer et al. 2008)

One species that that could be impacted by development is the serval (*Leptailurus serval*). The serval is a medium-sized carnivore that feeds primarily on rodents (Ramesh and Downs 2015), and is dependent on wetland habitats (Ramesh and Downs 2015/2) that are being rapidly lost globally (Dixon et al. 2016). The species is listed as Least Concern on the global IUCN Red List of threatened species (Thiel 2015), but is considered Near Threatened in South Africa (Friedmann and Daly 2004). Serval have declined throughout their range (Ramesh and Downs 2013), and the principal threats to the species are loss and degradation of their wetland habitat (Thiel 2011), trade of their skins (Kingdon and Hoffmann 2012), and persecution in response to perceived predation of poultry (Henley 1997), although they only rarely prey on livestock (Thiel 2015). Like many other felids, serval maintain stable home ranges where males typically have larger ranges than females (Sunquist and Sunquist 2002; Ramesh et al. 2015). While various factors (e.g. resource availability and physical attributes; Kie et al. 2002) affect carnivore home range size, in serval the availability of wetland habitats seems to be a key factor (Bowland 1990). Data on species ecology are critical to planning wildlife management and implementing conservation initiatives (Barrows et al. 2005), but there have been few studies on serval ecology, and conservation initiatives are hindered by poor knowledge of abundance (Ramesh and Downs 2013).

In this study, we firstly aimed to estimate the population density of servals at the Secunda Synfuels Operations plant, an industrial site in Mpumalanga province, South Africa, that includes a natural wetland within its boundaries (Fig. 1). We also aimed to assess the structure of this serval population, in order to make inferences about population dynamics.

**Fig. 1.**
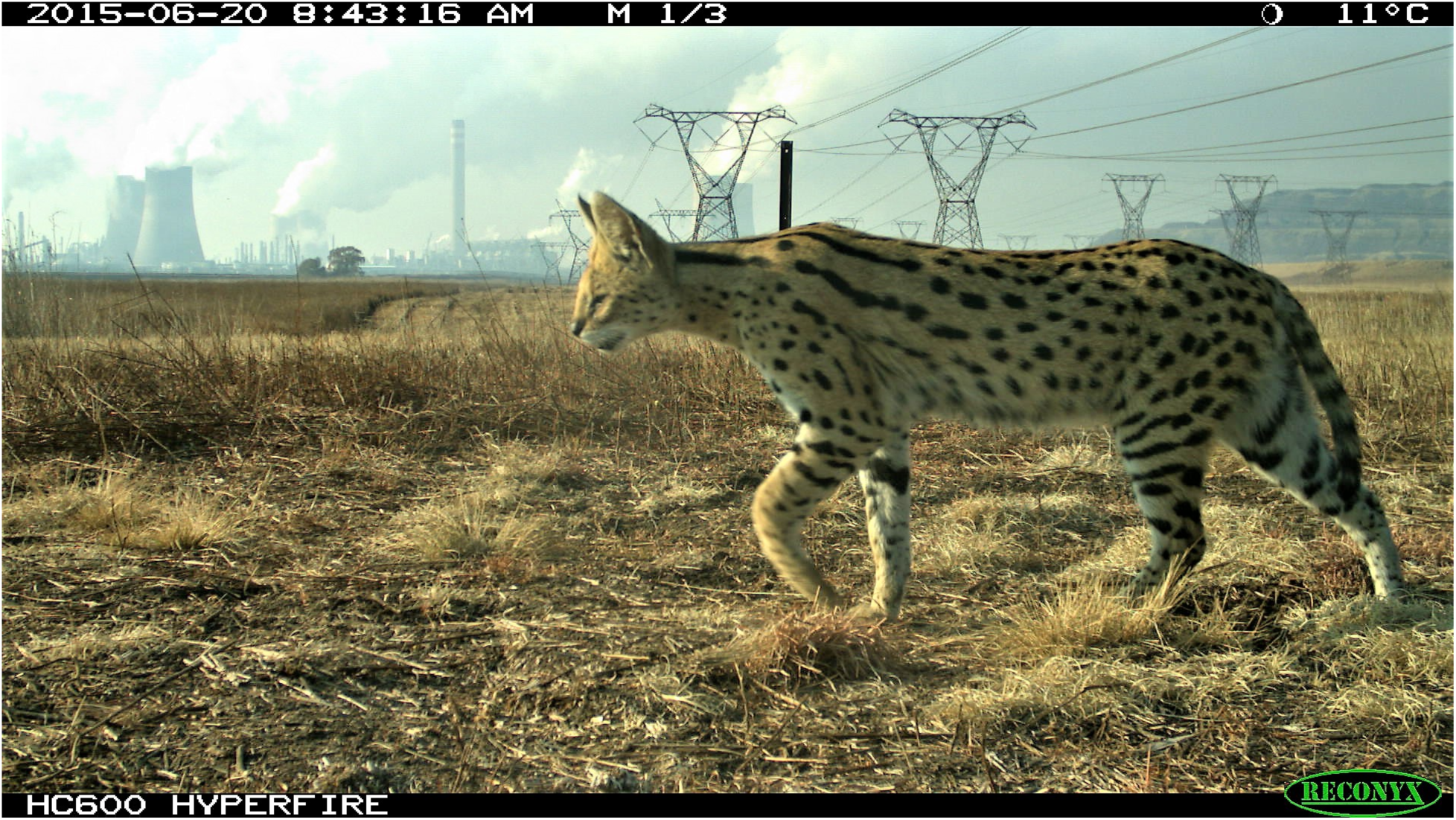
Camera trap image of a serval at the heavily industrialised Secunda Synfuels Operations plant in South Africa, recorded by Reconyx Hyperfire HC600 camera.

## 2 Materials and Methods

### 2.1 Ethics statement

This project is registered at the Animal Care and Use Committee of the University of Pretoria (Ethical clearance number: EC040-14 and V101-17) and the Mpumalanga Tourism and Parks Agency (Permit number: 5467 and 7282).

### 2.2 Study Area

The Secunda Synfuels Operations plant (hereafter referred to as Secunda) is a division of Sasol South Africa (PTY) Ltd, and is located in Secunda, Mpumalanga province, South Africa (Fig. 2). It consists of a primary area (a petrochemical plant) and a secondary area (which is made up of surrounding natural and disturbed vegetation). The secondary area (hereafter referred to as the study site) of Secunda Synfuels Operations covers an area of 79.4 km^2^ (central coordinates 26°31’45.62” S, 29°10’31.55” E). The secondary area is a gently to moderately undulating landscape on the Highveld plateau, supporting short to medium-high, dense, tufted grasses at different levels of disturbance. In places, small scattered wetlands (both man-made and natural), narrow stream alluvia, and occasional ridges or rocky outcrops interrupt the continuous grassland cover. Much of the study site (38%) is classified as relatively untransformed habitat, which is managed in accordance to Secunda Synfuels Operations Biodiversity Management Plan to conserve the natural areas from degradation and improve the ecological functionality of the disturbed land. The vegetation type is classified as Soweto Highveld Grassland (Rutherford et al. 2006), and the area falls into the Grassveld Biome (Mucina and Rutherford 2006). We used satellite images (Google 2014) to digitise the boundaries of four major habitat types (Disturbed, Grassland, Grass & wetland, and Wetland), which we used as site covariates in subsequent analyses.

**Fig. 2.**
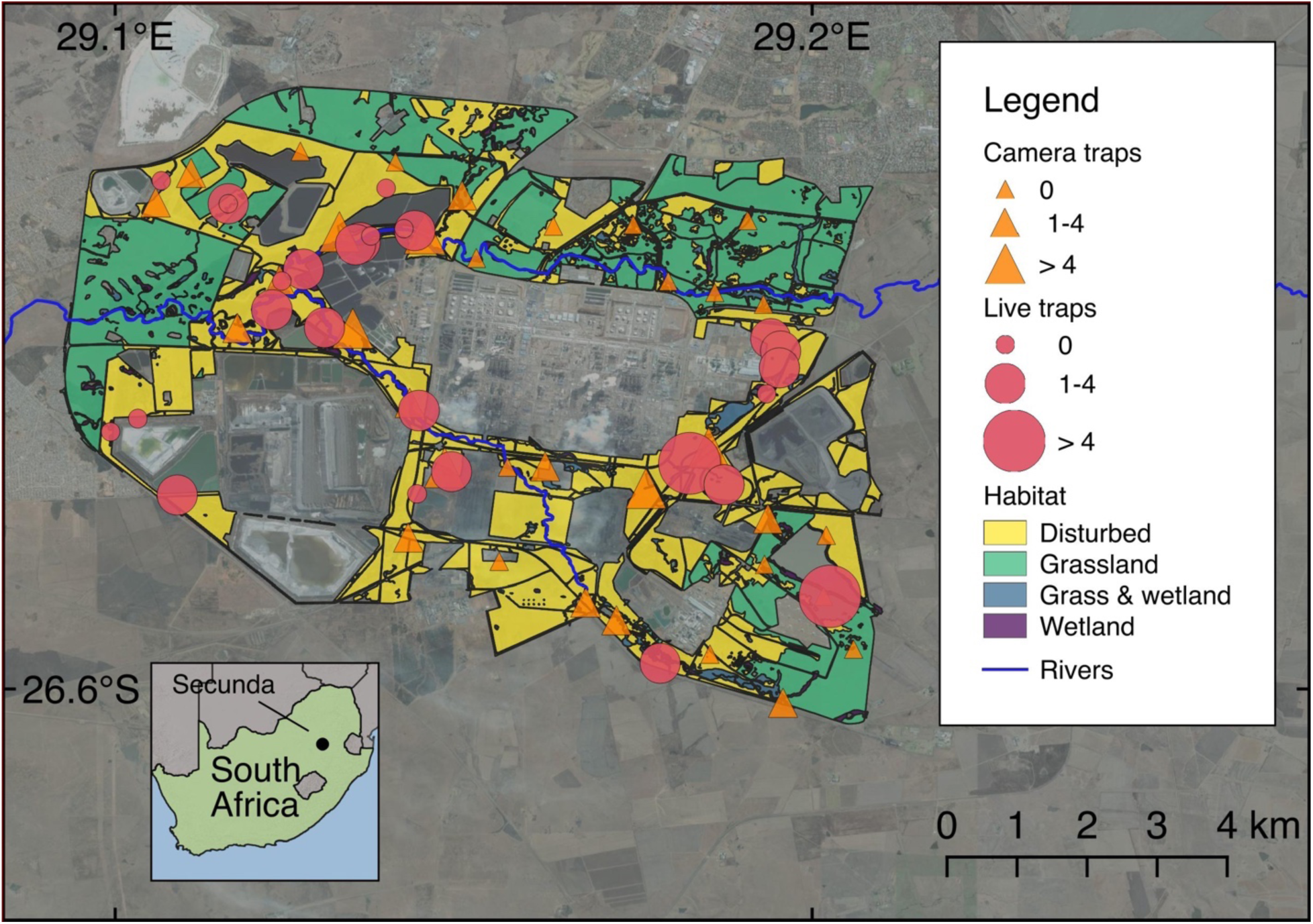
Map showing the locations of camera traps and live traps at the Secunda Synfuels Operations plant in South Africa. The size of points representing camera traps and live traps diameter is proportional to the number of individual serval captured. Major habitat types are also shown, along with satellite images illustrating the human-modified landscapes. Wetland and Grass & wetland habitat types are difficult to visualise at this scale as they occur in very close proximity to rivers.

The relatively unspoiled grassland represents the best form of Soweto Highveld Grassland on site. The characteristic species include *Cymbopogon pospischilii, Pollichia campestris, Walafrida densiflora, Eragrostis chloromelas, Gomphrena celosioides, Craibia affinis* and *Cineraria cf. savifraga* (Matthews 2016). The grassland habitat has a low basal cover due to grazing and during the rainy season the grass phytomass averages around 3-4 tons per hectare (de Wet 2016). The grass and wetland habitat occurs mostly within the transition zones or dry floodplains not typical of either wetland habitat or grassland habitat. These areas have a medium cover, and include some species typical of wetlands. The wetland habitat is dominated by species indicative of wetland zones and moist soils (Linström 2012). The phytomass here can be in excess of 5 tonnes per hectare, and the growth is up to 1.5 meters above the ground level. The disturbed habitat is dominated by weedy forbs with medium to very high density. The impact of the weedy forbs is a thicket of basal cover on the surface and up to 1.5 meters above ground level.

### 2.3 Camera trapping

The study was underpinned by a spatially explicit capture-recapture (SECR) framework. For SECR studies it is recommended that the camera trapping polygon be larger than the male home range size of the target species (Tobler and Powell 2013). The largest home range recorded for serval in South Africa (measured using minimum convex polygon) was 31.5 km^2^ (Bowland 1990). We first subdivided the study area in 34 grid cells measuring 1.2 km x 1.2 km (roughly equivalent to the size of smallest recorded serval home range (Ramesh and Downs 2015)). We then established an array of Reconyx Hyperfire HC600 camera traps at 34 camera trap stations (one in each grid cell) over an area of 79.4 km^2^ throughout the study site (Fig. 2). Mean spacing between camera traps was 1.2 km, and we placed camera traps on game trails and roads to maximise the probability of photographing servals, and to facilitate access for camera maintenance. We mounted camera traps on fence posts, 50 cm above the ground and 1 to 2 m from the trail. Vegetation in front of the camera traps was cleared to reduce false triggers.

We conducted three surveys from 2014 to 2015, with each survey running for 40 days (see Table 1 for dates). Camera traps were programmed to operate 24 hours per day, with a one minute delay between detections. We regarded each 24 hours as an independent sample. Camera trap positions were kept constant within each survey and between surveys. We visited each camera trap on a weekly basis to download the images, change batteries, and ensure the cameras remained in working order. Camera Base 1.4 (Tobler 2010) was used to catalogued the camera trap images. Since one of the assumptions of SECR models is that individuals are correctly identified, three authors (DL, WM, KE) identified individual serval in triplicate using distinct individual markings such as spot patterns and scars.

**Table 1.**
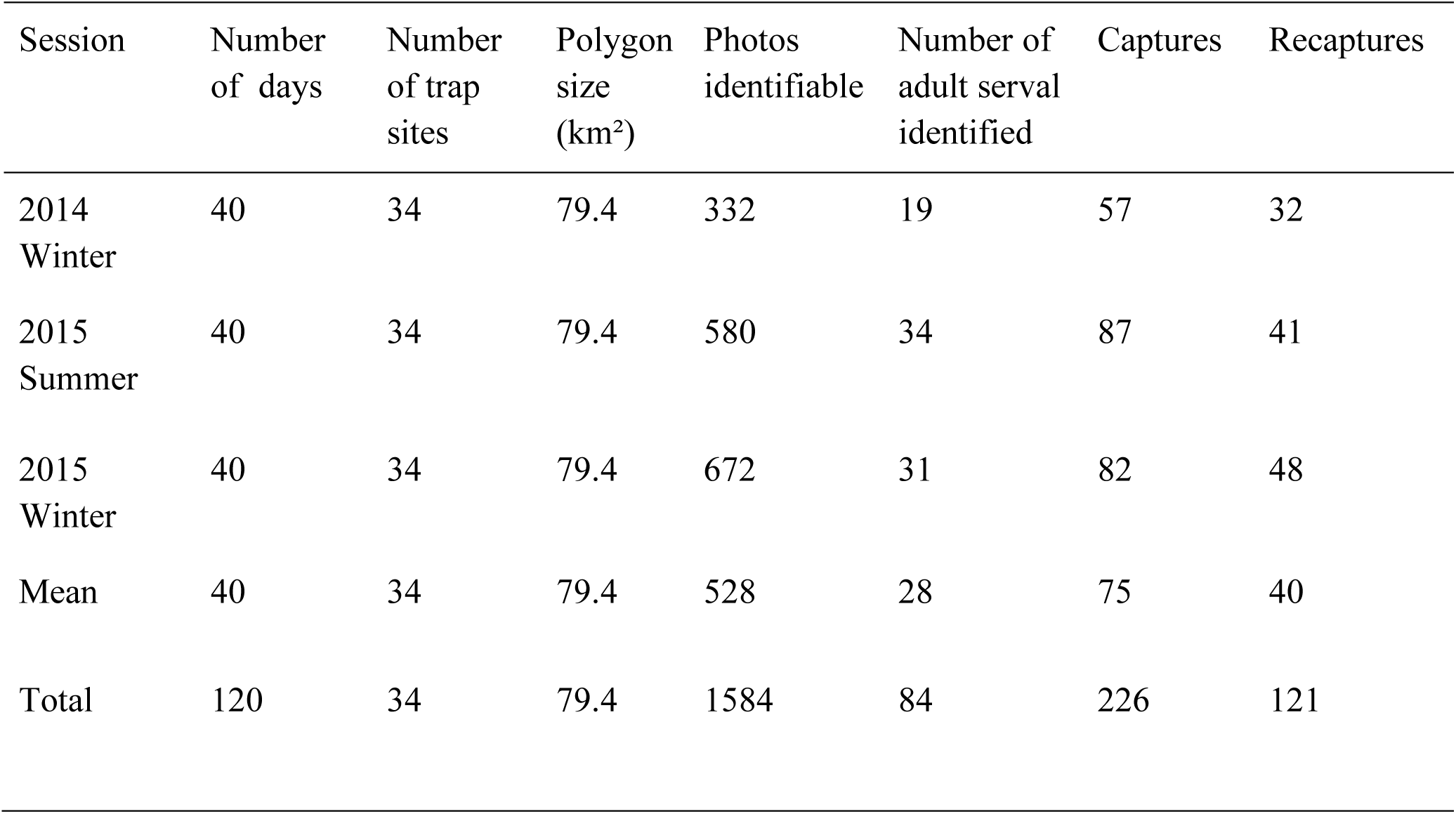
Summary of camera trapping effort at the study site during the winter of 2014, summer of 2015 and winter of 2015.

| Session | Number of days | Number of trap sites | Polygon size (km<sup>2</sup>) | Photos identifiable | Number of adult serval identified | Captures | Recaptures |
| --- | --- | --- | --- | --- | --- | --- | --- |
| 2014 Winter | 40 | 34 | 79.4 | 332 | 19 | 57 | 32 |
| 2015 Summer | 40 | 34 | 79.4 | 580 | 34 | 87 | 41 |
| 2015 Winter | 40 | 34 | 79.4 | 672 | 31 | 82 | 48 |
| Mean | 40 | 34 | 79.4 | 528 | 28 | 75 | 40 |
| Total | 120 | 34 | 79.4 | 1584 | 84 | 226 | 121 |

### 2.4 Live trapping

Live trapping formed part of a larger study investigating serval spatial and disease ecology. Due to low recapture success we did use these data to estimate densities. Rather, we used the live trapping data to estimate the capture rate and population structure of the serval population to validate our camera trapping study. Serval were trapped using 16 steel trap cages measuring 200 cm x 80 cm x 80 cm, deployed at 29 trap sites throughout the study site. Traps were baited with dead helmeted guineafowl (*Numida meleagris*) for a total of 287 trap nights between 2014 and 2017. Servals were immobilised by a veterinarian using one of the following drug combinations, as part of a study into optimising immobilisation protocols (Blignaut et al. in review): 1) KBM-5: ketamine (5.0 mg kg^−1^), butorphanol (0.2 mg kg^−1^), and medetomidine (0.08 mg kg^−1^); 2) KBM-8: ketamine (8.0 mg kg^−1^), butorphanol (0.2 mg kg^−1^), and medetomidine (0.08 mg kg^−1^); 3) ZM: zoletil (5.0 mg kg^−1^) and medetomidine (0.065 mg kg^−1^); 4) AM: alfaxalone (0.5 mg kg^−1^) and medetomidine (0.05 mg kg^−1^); or 5) ABM: alfaxalone (2.0 mg kg^−1^), butorphanol (0.2 mg kg^−1^), and medetomidine (0.08 mg kg^−1^). Drugs were administered intramuscularly using a blowpipe. If serval showed signs of inadequate drug dosages, they were topped-up with the same combinations. Where administered, medetomidine and butorphanol were pharmacologically antagonised with atipamezole (5 mg mg^−1^ medetomidine) and naltrexone (2 mg mg^−1^ butorphanol), respectively. After examination, animals were released at the same site where they were captured.

Animals with a mass of 3-8 kg were considered to be juveniles (up to approximately six months old, to the stage where the canines are developed). Servals with a mass of 8-11 kg were categorised as sub-adults (6-12 months old, just before they are sexually mature). Animals 11-15 kg (approximately 12 to 18 months and older) were considered to be adults (Sunquist and Sunquist 2002).

### 2.5 Data analysis

We estimated serval density by fitting likelihood based SECR (Efford 2004) models to camera trap data using the package secr (Efford 2017) in R version 3.4.3 (R Development Core Team 2017). The advantage of SECR models over traditional density estimation methods is that they do not require the use of subjective effective trapping areas, and instead estimate density directly (Tobler and Powell 2013). This is achieved by estimating the potential animal activity centres in a predefined area using spatial location data from the camera traps (Efford 2004). The spacing of the activity centres is related to the home range size of the animals, and as such the detection probability of each animal is a function of the distance from the camera trap to the activity centre. A key assumption of SECR models is that such activity centres are stationary for the period of study (closed population; Royle et al. 2015). Since serval are long lived animals exhibiting territoriality and we had relatively short survey period we believe that our study did not violate this assumption (van Aarde et al. 1986, Geertsema 1985).

Detection rate was modelled using a spatial detection function which is governed by two parameters; the encounter rate at the activity centre (detection probability;*λ*_0_) and a scale parameter (*σ*) which describes how the encounter rate declines with increased distance from the activity centre (Efford 2004). We tested for three different spatial detection functions since these might better model the utilisation distribution of the home range: half-normal, hazard and exponential. We ranked models based on Akaike information criterion corrected for small sample sizes (AICc), and found overwhelming support for the hazard rate spatial detection function (Table S3). All subsequent models were fitted with the hazard rate detection function.

We fitted SECR models by maximising the full likelihood where the scale parameter was kept constant, but we let the encounter rate vary by biologically plausible hypotheses to deal with heterogeneity in detection. The scale parameter is largely affected by home range size, and hence the sex of the animal (Sollmann et al. 2011/3). However, we were unable to determine the sex of individual serval from the photographs, and could therefore not model variation in the scale parameter due to sex. We first fitted a model in which we allowed the scale parameter to vary by year and season. This is because we expected that movement might be constrained in the wet season due to increased food resources (Courbin et al. 2013). We then fitted a model in which serval showed a behavioural response at *λ*_0_, as animals can become trap happy or trap shy (Wegge et al. 2004). Thirdly, we tested the effect of habitat on *λ*_0_, as serval prefer wetlands (Bowland 1990), which would result in higher detections in these habitats. We captured camera-specific habitat variables from the vegetation classification. Fourth, we coded each year and season as a separate session, and used the multi-session framework in secr to test the effect of season on serval density, with constant *λ*_0_. We lastly fitted a model in which *λ*_0_varied with both season and habitat type. These models were contrasted against a null model, in which all variables were kept constant.

We used AICc to rank models, considering models with ΔAICc < 2 to have equal support. We applied model averaging to the top models with equal support to reduce uncertainty (Burnham and Anderson 2004). The buffer width for analysis was set at 3,000 m, which resulted in the inclusion of an informal housing settlement and a residential area in the state space buffer. Since it is highly unlikely that serval will utilise these areas (as well as the primary industrial area), we excluded these areas (constituting approximately 25% of the area of the buffer) from the state space buffer (Fig. S1). All data and R code used for analysis are available in (Loock et al. 2018).

## 3 Results

### 3.1 Camera trapping

During a camera trapping effort of 3,590 trap days, we photographed a total 61 unique servals spanning three separate sessions (Table 1). The number of individual serval captures did not differ greatly between sessions, although the highest number was captured during the wet season of 2015 (Table 1, Fig. S2).

The two most parsimonious SECR models (ΔAICc < 2) both indicated that the encounter rate (*λ*_0_) was affected by habitat type (Table S3). While there was some support for serval density being session dependant (ΔAICc = 0.098; *AICc w =* 0.487; Table S3), there was also support for no effect of session (*AICc w* = 0.471, Table S3). To estimate serval density we therefore averaged the two most parsimonious models (ΔAICc < 2). Serval population density estimates at the study site varied from 76.20 (SE=22.22) to 101.21 (SE=20.66) animals per 100 km^2^ (Fig. 3a). Highest estimates were recorded during the dry seasons (Winter 2014: 101.21 [SE=20.66] & Winter 2015: 97.38 [SE=18.71]) compared to the single summer season (Summer 2015: 76.20 [SE=22.21]; Fig. 3a). Vegetation type had a significant effect on serval encounter rates, where grassland had the lowest encounter rate (0.04 [SE=0.01]) compared to wetlands with the highest (0.19 [SE=0.03]; Fig. 3b).

**Fig. 3.**
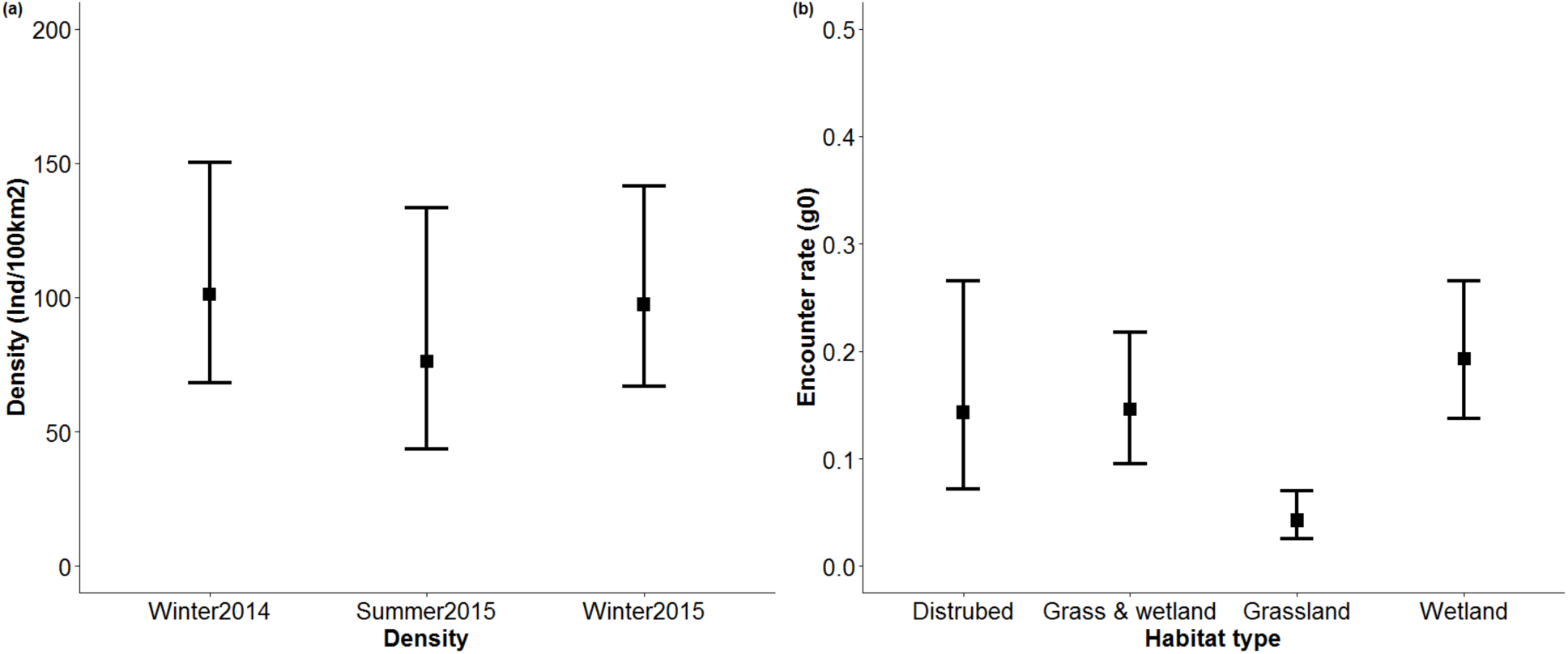
Serval density estimates for each camera trap survey conducted at the study site indicating a) influence of season on density, and b) effect of habitat type on serval encounter rate. Error bars represent asymmetric 95% confidence intervals.

### 3.2 Live trapping

We captured 65 individuals, of which four were also recaptured on a second occasion. This comprised of a total of 26 adult males, 19 adult females, 11 sub-adults, and seven juvenile animals. This resulted in a mean trapping success rate of 0.21 captures per trap night (excluding recaptures). Trapping success rate varied little between sessions (Fig. 4).

**Fig. 4.**
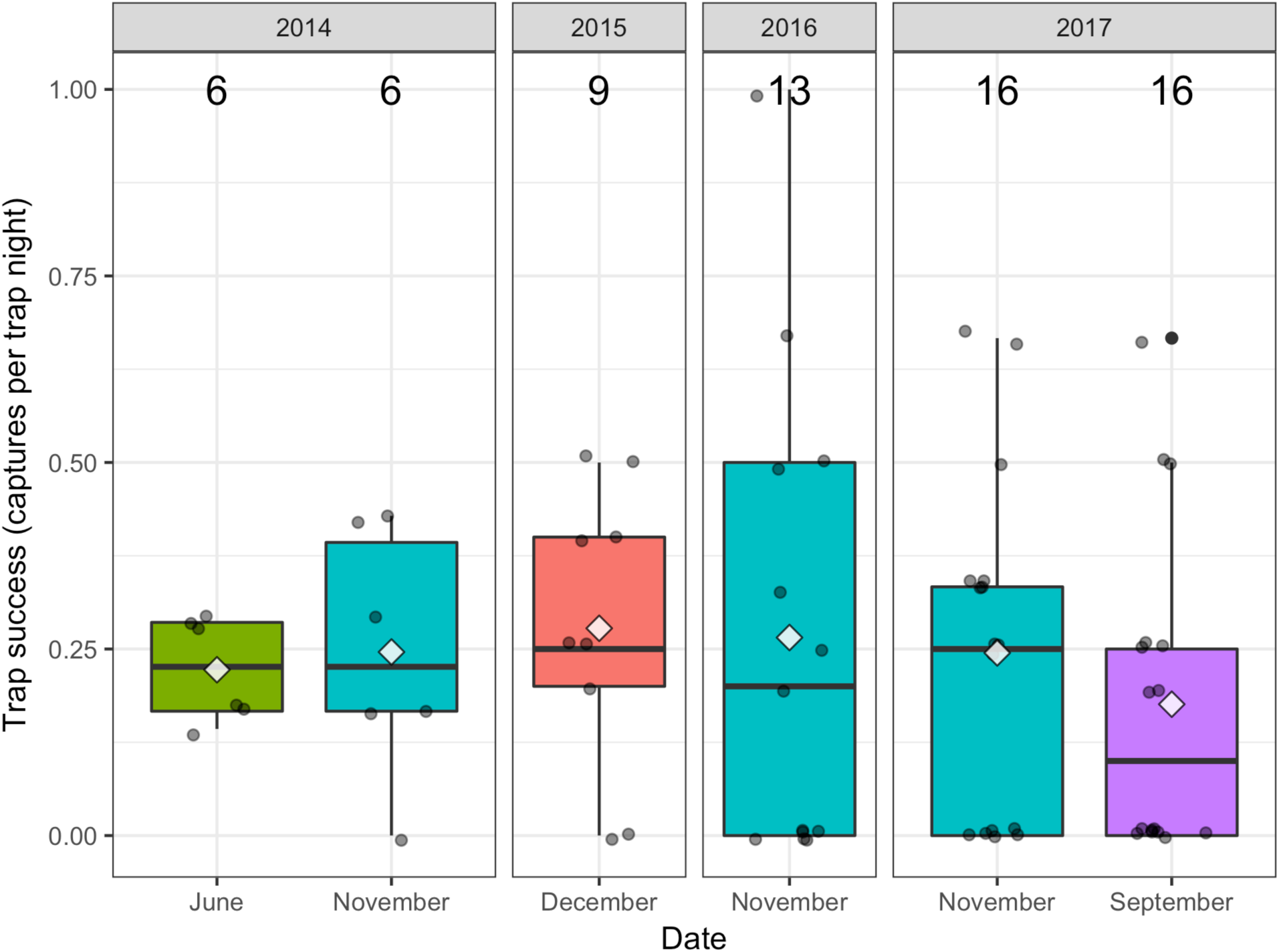
Box plot showing trap success rate for serval captures at the study site from 2014 to 2017. The middle bars represent the median value, white diamonds represent means, the top and bottom of the boxes represent the 75th and 25th percentiles respectively, the whiskers represent the maximum and minimum values, circles show the individual data points, and numbers give the sample size.

## 4 Discussion

### 4.1 Comparative serval density

In our three camera trap surveys at the study site at Secunda, we estimated serval population density to be 101.21, 76.20, and 97.38 animals per 100 km^2^, which are the highest densities recorded in the literature. Live trapping rates at Secunda were also extremely high (0.21 captures per trap night). Although there are no data available for serval live trapping rates in the literature, rates of 0.0015-0.017 captures per trap night are much more typical for other mesocarnivores such as jaguarundi (*Puma yagouaroundi*), oncilla (*Leopardus tigrinus*), tayra (*Eira barbara*), and feral cat (*Felis silvestris catus*) using cage traps (Molsher 2001; Michalski et al. 2007; McGregor, H W, Hampton J O, Lisle, D, Legge, S 2016), which are an order of magnitude lower than serval live capture rates at Secunda. Although great care must be taken when comparing trapping rates between different locations and species, the live trap rates at Secunda nevertheless appear to be consistently very high, which supports the high population densities estimated using camera trap data.

Our high estimates of serval densities at Secunda contrast with more typical densities reported in Luambe National Park in Zambia (9.9 animals per 100 km^2^ (Thiel 2011), Bwindi Impenetrable National Park in Uganda (9 animals per 100 km^2^ (Andama (2000), cited in Kingdon and Hoffmann 2012)), and on farmland in the Drakensberg Midlands, South Africa (6.5 animals per 100 km^2^ (Ramesh and Downs 2013)). However, there is evidence that serval can attain such high densities. For example, (Geertsema 1985) reported a serval density of 41.66 animals per 100 km^2^ in the Ngorongoro Crater, Tanzania.

High population densities of other carnivore species have also been reported in human-modified habitats such as urban areas. Coyotes (*Canis latrans*), raccoons (*Procyon lotor*), red foxes (*Vulpes vulpes*), and Eurasian badgers (*Meles meles*), for example, all thrive in urban landscapes (Bateman and Fleming 2012; Scott et al. 2014). Carnivore species able to adapt to urban environments often succeed in these areas due to high food availability, favourable climatic effects, and the reduced threat of intraguild predation because of the absence of larger apex predators (Fuller et al. 2010). We provide several, not necessarily mutually exclusive theories, to explain the high serval density we observed at Secunda.

Firstly, servals in the Secunda are protected from persecution. Such persecution can have large effects on carnivore densities. For example leopards (*Panthera pardus*) in livestock/game farming areas only attain around 20% of their potential density compared to protected areas free from persecution (Balme et al. 2010). Servals outside protected areas are frequently persecuted by livestock farmers (Henley 1997) as they are often mistakenly blamed for livestock predation (Skinner and Chimimba 2005), but at Secunda this is not the case, which could lead to higher population densities (Cardillo et al. 2004). Secondly, servals are the largest remaining carnivore species occurring at ecologically effective densities at Secunda, so there is little interspecific competition from larger carnivores. In other areas, the presence of other medium- and large-bodied carnivores could otherwise limit serval population densities (through intraguild predation), so their absence can lead to mesopredator release, such as through increased survival of young (Ritchie and Johnson 2009). For example, the absence of large carnivores such as lions (*Panthera leo*) and spotted hyaenas (*Crocuta crocuta*) in northern South Africa is thought to have led to the competitive release of cheetahs (*Acinonyx jubatus*) (Marnewick et al. 2007). Thirdly, the abundance of modified habitat at Secunda could also facilitate high serval population density. Disturbed habitat can be highly productive (Williams et al. 2018), and provide shelter and food resources for species such as rodents that serval prey upon (Taylor 2013), providing abundant food and in turn supporting a high abundance of serval.

Although the population density of serval recorded at Secunda was exceptionally high, the structure of this serval population was similar to those at other sites. The number of adult males per 100 adult females captured in live traps at Secunda was 137, which is within the range reported in the literature (50-220 in KwaZulu-Natal (Bowland 1990; Ramesh et al. 2016); 100 in the Ngorongoro Crater, Tanzania (Geertsema 1985)). Similarly, the proportion of the population at Secunda that was comprised of juvenile and sub-adult individuals (0.69) was very similar to other populations (0.64 in the Ngorongoro crater; (Geertsema 1985)). It therefore appears that although the serval population density at Secunda is very high, the structure of the population is not unusual, which is not indicative of a rapidly declining or increasing population size (Harris et al. 2008), supporting our findings that the serval population density at Secunda appears to be relatively stable. Although servals appear to thrive in close proximity to such a heavily industrialised site, we suggest that further research is conducted to identify any potential effects of industrial activity (Raiter et al. 2014), such as the influence of noise and air pollution on the physiology and behaviour of wildlife in the vicinity (Morris-Drake et al. 2017).

While we aimed to apply robust modelling, we address some caveats form our dataset. First, we were not able to include sex as a covariate in the SECR models, which could affect the density estimates. Simulation models suggest that excluding sex covariates can cause a negative bias in density estimates, thus overestimating density (Tobler and Powell 2013). As such it seems that estimates derived here can be regarded as optimistic. Nonetheless, the scale parameter used in the models (sigma = 268 m) falls within range of observed daily movements of serval elsewhere (538m (Perrin 2002); 0-500m (van Aarde et al. 1986)). This suggests that the estimated 95% confidence interval should encompass the true estimates, albeit on the lower side of interval. Secondly, the placement of the camera traps was constrained by the vegetation conditions, in order to enable access to camera traps by foot or by vehicle. This could have introduced sampling bias as traps were not placed at random in relation to activity centres. However, maximising the detection of individuals in order to obtain adequate samples outweighs the potential bias caused by biased trap placement (Tobler and Powell 2013). Finally, there might be concern regarding population closure since our trapping period spanned 40 days and we had a high percentage of single detections. We highlight that SECR models appear to be robust against transience (Royle et al. 2015) and that longer surveys tend to yield more robust estimates than short periods (Jędrzejewski et al. 2016).

### 4.2 The impacts of modified landscapes

In recent years the expansion of infrastructure has progressed more rapidly than during any other period in history (Laurance et al. 2015), and industrial sites such as mines and fossil fuel processing plants are not the only developments that could have impacts on wildlife. The growing road network, for example (Ibisch et al. 2016), has large direct and indirect ecological impacts such as causing wildlife-vehicle collisions, polluting the environment, disrupting animal migrations and gene flow, and providing access to invading species and humans, facilitating further degradation (Laurance et al. 2009; Sloan et al. 2016). The rapidly growing number of hydroelectric dams (Zarfl et al. 2014) increases the risk of habitat fragmentation through deforestation, in addition to disrupting freshwater ecosystems (Finer and Jenkins 2012). Similarly, the development of urban and agricultural areas fragments and destroys habitats (Ripple et al. 2014). Consequently, delineating how the changing environment affects biodiversity will be an increasingly important theme of future research.

But not all the impacts of anthropogenic development on wildlife are negative. The high serval densities at Secunda are remarkable as the site is very heavily industrialised. Nature reserves and exclusion zones surrounding industrialised areas such as Secunda have the potential to balance resource utilisation with biodiversity conservation (Edwards et al. 2014). Some industrial installations such as mines have created nature reserves, which can benefit biodiversity conservation. The Mbalam iron ore mine in Cameroon has set aside land to protect rare forest mammals (Edwards et al. 2014). Private nature reserves created around the Venetia diamond mine in South Africa and the Jwaneng diamond mine in Botswana support a broad complement of large mammals including elephants (*Loxodonta africana*), lions (*Panthera leo*), leopards (*Panthera pardus*), cheetahs, African wild dogs (*Lycaon pictus*), brown hyaenas (*Hyaena brunnea*), and black-backed jackals (*Canis mesomelas*) (Smallie and O’connor 2000; Kamler et al. 2007; Houser et al. 2009; Jackson et al. 2014). The Sperrgebiet exclusion zone in Namibia, established to protect diamond deposits (Edwards et al. 2014), has now been proclaimed a National Park (Wiesel 2010). The consequent changes in the ecological functions of these human modified areas can produce a new combination of species, sometimes modifying and, in some cases, increasing the local richness (Hobbs et al. 2006; Pautasso et al. 2011).

Studies such as this highlight the complexity of the relationship between wildlife and the human-modified environment, and suggest that the potential conservation value of industrialised sites should not be overlooked. This underscores the importance of sound ecological management in these areas. Such sites could be incorporated into wildlife management plans, and could help to achieve goals such as the conservation of threatened species. This could be achieved, for example, through the formation of partnerships between industry and the non-profit sector or governmental agencies, such as the partnership between Eskom and the Endangered Wildlife Trust (EWT) to reduce the threats posed by electricity infrastructure to wildlife in South Africa (Jenkins et al. 2010).

## 5 Conclusion

Servals occur at much greater densities at Secunda than have been recorded elsewhere. Capture rates on both camera traps and live traps were remarkably high. High densities may be due to favourable conditions such as a high abundance of rodent prey and the absence of persecution or competitor species. Despite the highly industrialised nature of the site, serval population structure appears to be similar to other natural sites. We suggest that the potential value of industrial sites, where they include areas of relatively natural habitats, may be underappreciated by conservationists, and that these sites could help meet conservation objectives.

## Competing interests

DJEL author is a full-time employee of Secunda Synfuels Operations (a division of Sasol South Africa (Pty) Ltd) as Land & Biodiversity Manager. Secunda Synfuels Operations had no role in study design, data collection and analysis, decision to publish, or preparation of the manuscript. STW, KWE, WSM, and LHS declare that they have no potential competing interests.

## Acknowledgements

We would like to thank Secunda Synfuels Operations > Div of Sasol South Africa (Pty) Ltd. for supporting this research, the Faculty of Natural and Agricultural Science, University of the Free State, and the Wildlife Resource Association (WRA). STW was supported by a postdoctoral grant from the University of Venda. LHS was supported by the National Research Foundation of South Africa (Grant Nr: 107099) and the University of Venda. We are grateful to Chris Sutherland for advice on SECR modelling. Finally, we would like to thank Matt Hayward and an anonymous reviewer for their suggestions, which helped to improve the manuscript.

## Author Biographies

Daan Loock is a Land & Biodiversity Manager at Secunda Synfuels Operations > Div of Sasol South Africa (Pty) Ltd. He is currently involved in biodiversity studies and especially a serval research program for the last couple of years. Address: Secunda Synfuels Operations, Private Bag X1000, Secunda, 2302, South Africa.

Samual T. Williams is a Postdoctoral Research Fellow in the Department of Zoology at the University of Venda. His research centres around the theme of carnivores in the Anthropocene, including carnivore conservation, road ecology, and ecosystem services. Address: Department of Zoology, School of Mathematical & Natural Sciences, University of Venda, Private Bag X5050, Thohoyandou 0950, South Africa.

Kevin W. Emslie is an MSc. candidate in the Department of Zoology at the University of Venda. His research focuses on the impacts of human-modified landscapes on small- and mesocarnivore ecology. Address: Department of Zoology, School of Mathematical & Natural Sciences, University of Venda, Private Bag X5050, Thohoyandou 0950, South Africa.

Wayne S. Matthews has completed his MSc and PhD, which dealt with the vegetation and ecology of the North-eastern Mountain Sourveld, and ecology with dynamics of Sand Forest of Maputaland. Address: Wildlife Resource Association, PO Box 1288, Umhlali, 4390, South Africa.

Lourens H. Swanepoel is a Senior Lecturer in Conservation Biology at the Department of Zoology, University of Venda. His research interests include carnivore conservation, human carnivore conflict and ecosystem services. Address: Department of Zoology, School of Mathematical & Natural Sciences, University of Venda, Private Bag X5050, Thohoyandou 0950, South Africa.

## Supplementary information

**Fig. S1.**
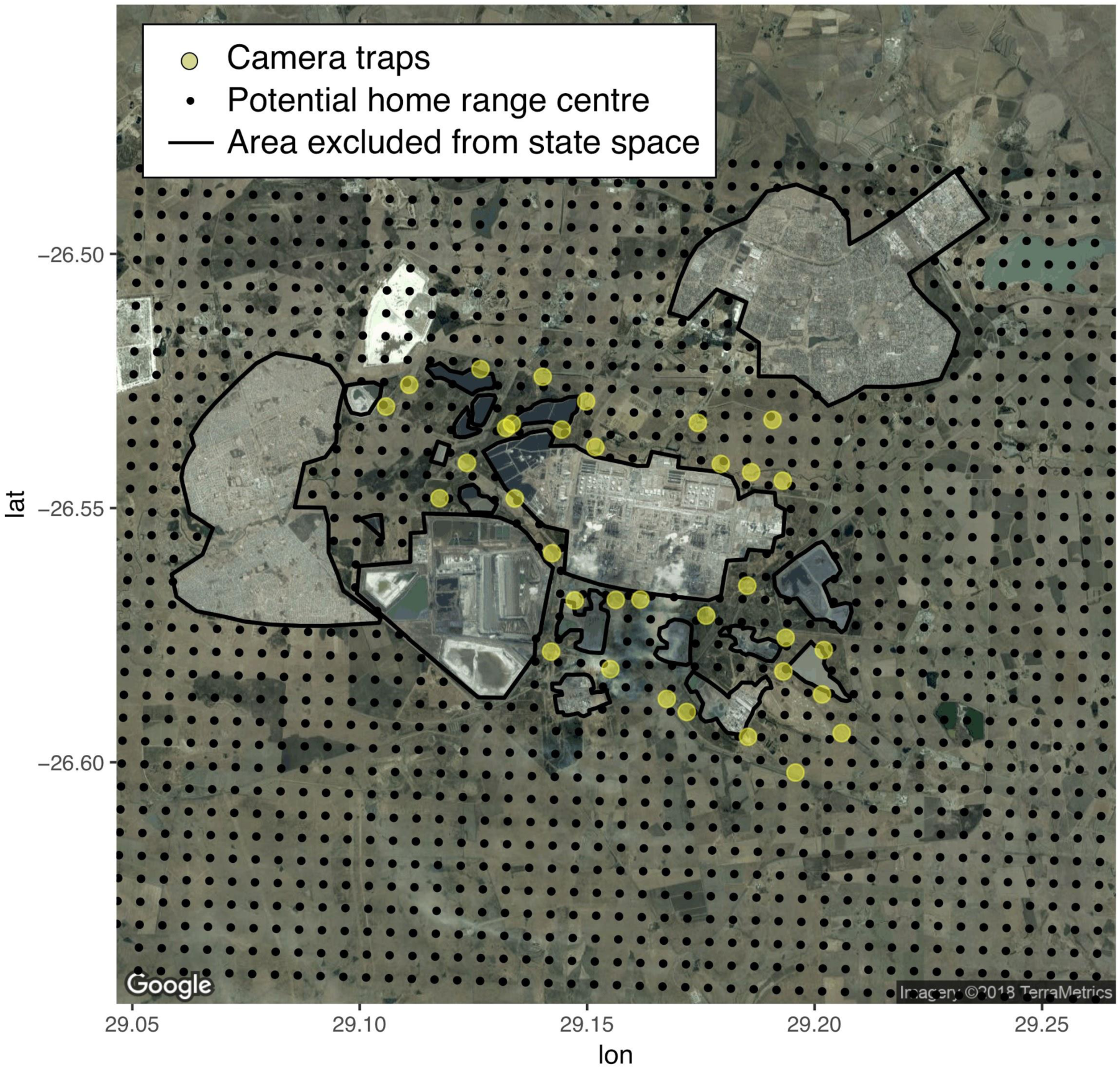
Map of Secunda in South Africa showing the locations of camera traps and potential home range centres, illustrating the areas excluded from the state space.

**Fig. S2.**
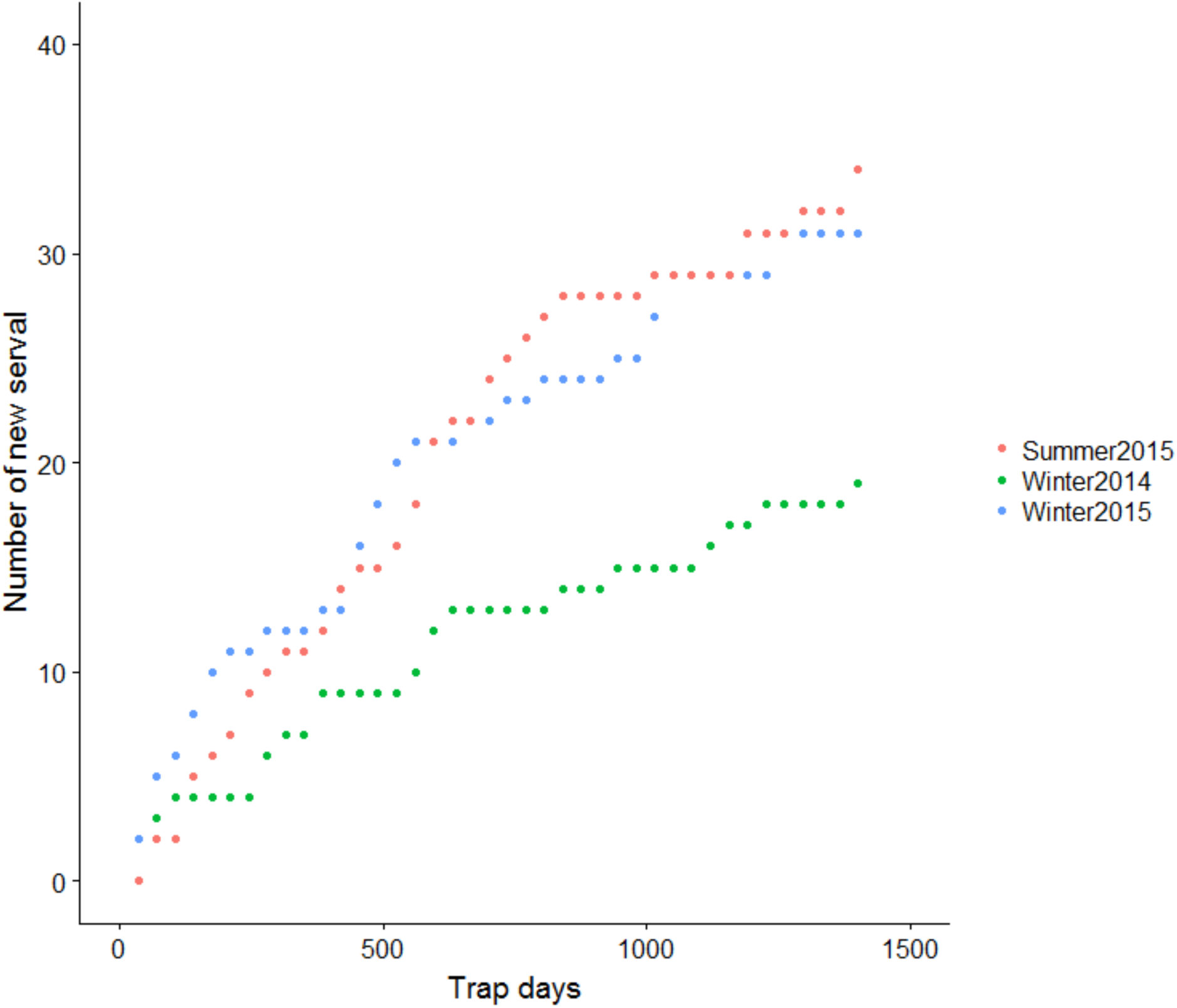
Cumulative frequency curve showing the relationship between the cumulative number of individual serval identified on the camera traps at Secunda in South Africa.

**Table S3.** Modelling results showing: a) (Mhab.sec) the effect of habitat type on detection probability (g0); b) (Mdens.habitat) the effect of session on density and habitat on detection probability; c) as model b, but allows the scale parameter (sigma) to vary with session; d) (Mb.sec) the effect of behavioural response on detection probability; e) (Mseason) the effect of density affected by season and year; and f) (m0.sec) the null model.

|  | model | detectfn | npar | logLik | AIC | AICc | dAICc | AICc wt |
| --- | --- | --- | --- | --- | --- | --- | --- | --- |
| Mhab.sec | D~1<br>g0~Habitat<br>sigma~1<br>z~1 | hazard<br>rate | 7 | -<br>1015.426<br>872 | 2044.8<br>5 | 2046.3<br>3 | 0.00 | 0.47 |
| Mdens.habitat | D~session<br>g0~Habitat<br>sigma~1<br>z~1 | hazard<br>rate | 9 | -<br>1012.996<br>398 | 2043.9<br>9 | 2046.4<br>3 | 0.10 | 0.45 |
| Mdens.habitat.ses | D~session<br>g0~Habitat<br>sigma~session<br>z~1 | hazard<br>rate | 11 | -<br>1012.124<br>715 | 2046.2<br>5 | 2049.9<br>2 | 3.59 | 0.08 |
| Mb.sec | D~1 g0~b<br>sigma~1<br>z~1 | hazard<br>rate | 5 | -<br>1028.514<br>939 | 2067.0<br>3 | 2067.8<br>0 | 21.47 | 0.00 |
| Mseason | D~session<br>g0~1<br>sigma~1<br>z~1 | hazard<br>rate | 6 | -<br>1030.089<br>274 | 2072.1<br>8 | 2073.2<br>7 | 26.94 | 0.00 |
| M0.sec | D~1 g0~1<br>sigma~1<br>z~1 | hazard<br>rate | 4 | -<br>1033.592<br>632 | 2075.1<br>9 | 2075.6<br>9 | 29.37 | 0.00 |

## References

van Aarde, R. J., and J. D. Skinner. 1986. Pattern of space use by relocated servals *Felis serval*. African Journal of Ecology 24: 97–101. doi:10.1111/j.1365-2028.1986.tb00348.x.

Balme, G. A., R. Slotow, and L. T. B. Hunter. 2010. Edge effects and the impact of non-protected areas in carnivore conservation: leopards in the Phinda-Mkhuze Complex, South Africa. Animal Conservation 13: 315–323. doi:10.1111/j.1469-1795.2009.00342.x.

Barbosa, A. M., D. Fontaneto, L. Marini, and M. Pautasso. 2010. Positive regional species–people correlations: a sampling artefact or a key issue for sustainable development? Animal Conservation 13: 446–447. doi:10.1111/j.1469-1795.2010.00402.x.

Barrows, C. W., M. B. Swartz, W. L. Hodges, M. F. Allen, J. T. Rotenberry, B.-L. Li, T. A. Scott, and X. Chen. 2005. A framework for monitoring multiple-species conservation plans. Journal of Wildlife diseases D9: 1333–1345. doi:10.2193/0022-541X(2005)69[1333:AFFMMC]2.0.CO;2.

Bateman, P. W., and P. A. Fleming. 2012. Big city life: carnivores in urban environments. Journal of Zoology 28: 1–23. doi:10.1111/j.1469-7998.2011.00887.x.

Blignaut, C., G. Steenkamp, J. Hewlett, D. Loock, R. Emslie, and G. E. Zeiler. In review. Preliminary findings of free-ranging serval (Leptailurus serval) chemical capture.

Bowland, J. M. 1990. Diet, home range and movement patterns of serval on farmland in Natal.

Cardillo, M., A. Purvis, W. Sechrest, J. L. Gittleman, J. Bielby, and G. M. Mace. 2004. Human population density and extinction risk in the world’s carnivores. PLoS biology 2: E197. doi:10.1371/journal.pbio.0020197.

Chapin, F. S., E. S. Zavaleta, V. T. Eviner, R. L. Naylor, P. M. Vitousek, H. L. Reynolds, D. U. Hooper, S. Lavorel, O. E. Sala, S. E. Hobbie, M. C. Mack, and S. Díaz. 2000. Consequences of changing biodiversity. Nature 405: 234–242. doi:10.1038/35012241.

Courbin, N., D. Fortin, and C. Dussault. 2013. Multi-trophic resource selection function enlightens the behavioural game between wolves and their prey. Journal of Animal Ecology: 82(5): 1062–1071. doi:10.1111/1365-2656.12093.

Dixon, M. J. R., J. Loh, N. C. Davidson, C. Beltrame, R. Freeman, and M. Walpole. 2016. Tracking global change in ecosystem area: The Wetland Extent Trends index. Biological Conservation 193: 27–35. doi:10.1016/j.biocon.2015.10.023.

Edwards, D. P., S. Sloan, L. Weng, P. Dirks, J. Sayer, and W. F. Laurance. 2014. Mining and the African Environment. Conservation Letters 7: 302–311. doi:10.1111/conl.12076.

Efford, M. 2004. Density estimation in live-trapping studies. Oikos 106: 598–610. doi:10.1111/j.0030-1299.2004.13043.x.

Efford, M. G. 2017. secr: Spatially explicit capture-recapture models. R package version 3.1.3. Available from https://cran.R-project.org/package=secr.

Ellis, E. C., K. Klein Goldewijk, S. Siebert, D. Deborah Lightman, and N. Ramankutty. 2010. Anthropogenic transformation of the biomes, 1700 to 2000. Global Ecology and Biogeography 19: 589–606. doi:10.1111/j.1466-8238.2010.00540.x/full.

Ellis, E. C., J. O. Kaplan, D. Q. Fuller, S. Vavrus, K. Klein Goldewijk, and P. H. Verburg. 2013. Used planet: a global history. Proceedings of the National Academy of Sciences 110: 7978–7985. doi:10.1073/pnas.1217241110.

Finer, M., and C. N. Jenkins. 2012. Proliferation of hydroelectric dams in the Andean Amazon and implications for Andes-Amazon connectivity. PLoS One 7: e35126. doi:10.1371/journal.pone.0035126.

Friedmann, Y., and B. Daly. 2004. Red Data Book of the Mammals of South Africa: a Conservation Assessment. Johannesburg: Endangered Wildlife Trust.

Fuller, T., S. Destefano, and P. S. Warren. 2010. Carnivore behavior, ecology and relationship to urbanization. In Urban Carnivores: Ecology, Conflict, and Conservation, ed. S. D. Gehrt, S. P. D. Riley, and B. L. Cypher, 13–20. Baltimore: Johns Hopkins University Press.

Geertsema, A. A. 1985. Aspects of the ecology of the serval *Leptailurus serval* in the Ngorongoro Crater, Tanzania. Netherlands Journal of Zoology 35: 527–610. doi:10.1163/002829685X00217.

Google. 2014. Satellite imagery. Sources: CNES/Airbus. Image date 01 May 2014. Accessed May 10, 2014. Available from https://www.google.co.za/maps

Harris, N. C., M. J. Kauffman, and L. S. Mills. 2008. Inferences about ungulate population dynamics derived from age ratios. Journal of Wildlife Management 72: 1143–1151. doi:10.2193/2007-277.

Henley, S. 1997. On the proposed reintroduction of serval (Felis serval) into the Great Fish River Reserve, Eastern Cape. Port Elizabeth: University of Port Elizabeth.

Hobbs, R. J., S. Arico, J. Aronson, J. S. Baron, P. Bridgewater, V. A. Cramer, P. R. Epstein, J. J. Ewel, et al. 2006. Novel ecosystems: theoretical and management aspects of the new ecological world order. Global Ecology and Biogeography 15: 1–7. doi:10.1111/j.1466-822X.2006.00212.x.

Houser, A., M. J. Somers, and L. K. Boast. 2009. Home range use of free-ranging cheetah on farm and conservation land in Botswana. South African Journal of Wildlife Research 39: 11–22. doi:10.3957/056.039.0102.

Ibisch, P. L., M. T. Hoffmann, S. Kreft, G. Pe’er, V. Kati, L. Biber-Freudenberger, D. A. DellaSala, M. M. Vale, P. R. Hobson, and N. Selva. 2016. A global map of roadless areas and their conservation status 354: 1423–1427. doi:10.1126/science.aaf7166.

Jackson, C. R., R. J. Power, R. J. Groom, E. H. Masenga, E. E. Mjingo, R. D. Fyumagwa, E. RØskaft, and H. Davies-Mostert. 2014. Heading for the hills: risk avoidance drives den site selection in African wild dogs. PLoS One 9: e99686. doi:10.1371/journal.pone.0099686.

Jenkins, A. R., J. J. Smallie, and M. Diamond. 2010. Avian collisions with power lines: a global review of causes and mitigation with a South African perspective. Bird Conservation International 20: 263–278. doi:10.1017/S0959270910000122.

Kamler, J. F., H. T. Davies-Mostert, L. Hunter, and D. W. Macdonald. 2007. Predation on black-backed jackals (*Canis mesomelas*) by African wild dogs (*Lycaon pictus*). African Journal of Ecology 45: 667–668. doi:10.1111/j.1365-2028.2007.00768.x.

Kie, J. G., R. Terry Bowyer, M. C. Nicholson, B. B. Boroski, and E. R. Loft. 2002. Landscape heterogeneity at differing scales: Effects on spatial distribution of mule deer. Ecology 83: 530–544. doi:10.2307/2680033.

Kingdon, J., and M. Hoffmann, eds. 2012. Mammals of Africa. New York: Bloomsbury.

Laurance, W. F., M. Goosem, and S. G. W. Laurance. 2009. Impacts of roads and linear clearings on tropical forests. Trends in Ecology & Evolution 24: 659–669. doi:10.1016/j.tree.2009.06.009.

Laurance, W. F., A. Peletier-Jellema, B. Geenen, H. Koster, P. Verweij, P. Van Dijck, T. E. Lovejoy, J. Schleicher, and M. van Kuijk. 2015. Reducing the global environmental impacts of rapid infrastructure expansion. Current Biology 25: R259–262. doi:10.1016/j.cub.2015.02.050.

Lindenmayer, D. B., J. Fischer, A. Felton, M. Crane, D. Michael, C. Macgregor, R. Montague-Drake, A. Manning, and R. J. Hobbs. 2008. Novel ecosystems resulting from landscape transformation create dilemmas for modern conservation practice. Conservation Letters 1: 129–135. doi:10.1111/j.1755-263x.2008.00021.x.

LinstrÖm, A. 2012. Sasol: Secunda Wetland Study. Wet Earth Eco-Specs, Lydenburg.

Loock, D., S. Williams, K. Emslie, M. Somers, W. S. Matthews, and L. Swanepoel. 2018. Serval (*Leptailurus serval*) camera trap and live trap dataset at Secunda, South Africa. Figshare. Accessed 22 March 2018. Available from https://figshare.com/s/bec3a5d725d8c11a5842. doi:10.6084/m9.figshare.5729124.

Marnewick, K., A. Beckhelling, D. Cilliers, E. Lane, G. Mills, K. Herring, P. Caldwell, R. Hall, and S. Meintjes. 2007. The status of the cheetah in South Africa. Cat News Special Issue 3 – Cheetahs in Southern Africa: 22–31.

Matthews, W. S. 2016. Baseline assessment of state of terrestrial flora for the “secondary area” of the SASOL Secunda site. Part of the state of biodiversity assessment – WetEarth. WSM Eco Services, Nelspruit.

McGregor, H. W., J. O. Hampton, D. Lisle, and S. Legge. 2016. Live-capture of feral cats using tracking dogs and darting, with comparisons to leg-hold trapping. Wildlife Research 43: 313–322. doi:10.1071/WR15134.

McKinney, M. L. 2002. Urbanization, biodiversity, and conservation: the impacts of urbanization on native species are poorly studied, but educating a highly urbanized human population about these impacts can greatly improve species conservation in all ecosystems. BioScience 52: 883–890. doi:10.1641/0006-3568(2002)052[0883:UBAC]2.0.CO;2.

Michalski, F., P. G. Crawshaw Jr, T. G. de Oliveira, and M. E. Fabián. 2007. Efficiency of box-traps and leg-hold traps with several bait types for capturing small carnivores (Mammalia) in a disturbed area of southeastern Brazil. Revista de Biologia Tropical 55: 315–320.

Molsher, R. L. 2001. Trapping and demographics of feral cats (*Felis catus*) in central New South Wales. Wildlife Research 28: 631–636. doi:10.1071/WR00027.

Morris-Drake, A., A. M. Bracken, and J. M. Kern. 2017. Anthropogenic noise alters dwarf mongoose responses to heterospecific alarm calls. Environmental pollution. 223: 476–483. doi:10.1016/j.envpol.2017.01.049.

Mucina, L., and M. C. Rutherford. 2006. The Vegetation of South Africa, Lesotho and Swaziland. Pretoria: South African National Biodiversity Institute.

Northrup, J. M., and G. Wittemyer. 2013. Characterising the impacts of emerging energy development on wildlife, with an eye towards mitigation. Ecology Letters 16: 112–125. doi:10.1111/ele.12009.

Pautasso, M., K. BÖhning-Gaese, P. Clergeau, V. R. Cueto, M. Dinetti, E. Fernández-Juricic, M.-L. Kaisanlahti-Jokimäki, J. Jokimäki, M. L. McKinney, N. S. Sodhi, D. Storch, L. Tomialojc, P. J. Weisberg, J. Woinarski, R. A. Fuller, and E. Cantarello. 2011. Global macroecology of bird assemblages in urbanized and semi-natural ecosystems. Global Ecology and Biogeography 20: 426–436. doi:10.1111/j.1466-8238.2010.00616.x.

Perrin, M. R. 2002. Space use by a reintroduced serval in Mount Currie Nature Reserve. South African Journal of Wildlife Research 32: 79–86.

Raiter, K. G., H. P. Possingham, S. M. Prober, and R. J. Hobbs. 2014. Under the radar: mitigating enigmatic ecological impacts. Trends in Ecology & Evolution 29: 635–644. doi:10.1016/j.tree.2014.09.003.

Ramesh, T., and C. T. Downs. 2015/2. Impact of land use on occupancy and abundance of terrestrial mammals in the Drakensberg Midlands, South Africa. Journal for Nature Conservation 23: 9–18. doi:10.1016/j.jnc.2014.12.001.

Ramesh, T., and C. T. Downs. 2013. Impact of farmland use on population density and activity patterns of serval in South Africa. Journal of Mammalogy 94: 1460–1470. doi:10.1644/13-MAMM-A-063.1.

Ramesh, T., and C. T. Downs. 2015. Diet of serval (*Leptailurus serval*) on farmlands in the Drakensberg Midlands, South Africa. Mammalia. doi:10.1515/mammalia-2014-0053.

Ramesh, T., R. Kalle, and C. T. Downs. 2015. Sex-specific indicators of landscape use by servals: Consequences of living in fragmented landscapes. Ecological Indicators 52: 8–15. doi:10.1016/j.ecolind.2014.11.021.

Ramesh, T., R. Kalle, and C. T. Downs. 2016. Spatiotemporal variation in resource selection of servals: insights from a landscape under heavy land-use transformation. Journal of Mammalogy 97: 554–567. doi:10.1093/jmammal/gyv201.

R Development Core Team. 2017. R: A language and environment for statistical computing. Version 3.4.3. R Foundation for statistical computing, Vienna. Available from https://www.R-project.org/

Ripple, W. J., J. A. Estes, R. L. Beschta, C. C. Wilmers, E. G. Ritchie, M. Hebblewhite, J. Berger, B. Elmhagen, M. Letnic, M. P. Nelson, O. J. Schmitz, D. W. Smith, A. D. Wallach, and A. J. Wirsing. 2014. Status and ecological effects of the world’s largest carnivores. Science 343: 1241484–1241484. doi:10.1126/science.1241484.

Ritchie, E. G., and C. N. Johnson. 2009. Predator interactions, mesopredator release and biodiversity conservation. Ecology Letters 12: 982–998. doi:10.1111/j.1461-0248.2009.01347.x.

Royle, J. A., J. Andrew Royle, A. K. Fuller, and C. Sutherland. 2015. Spatial capture–recapture models allowing Markovian transience or dispersal. Population Ecology 58: 53–62. doi:10.1007/s10144-015-0524-z.

Sax, D. F., and S. D. Gaines. 2003. Species diversity: from global decreases to local increases. Trends in Ecology & Evolution 18: 561–566. doi:10.1016/S0169-5347(03)00224-6.

Scott, D. M., M. J. Berg, B. A. Tolhurst, A. L. M. Chauvenet, G. C. Smith, K. Neaves, J. Lochhead, and P. J. Baker. 2014. Changes in the distribution of red foxes (*Vulpes vulpes*) in urban areas in Great Britain: findings and limitations of a media-driven nationwide survey. PLoS One 9: e99059. doi:10.1371/journal.pone.0099059.

Skinner, J. D., and C. T. Chimimba. 2005. The Mammals of the Southern African Sub-region. Cambridge: Cambridge University Press.

Sloan, S., B. Bertzky, and W. F. Laurance. 2016. African development corridors intersect key protected areas. African Journal of Ecology 55: 731–737. doi:10.1111/aje.12377.

Smallie, J. J., and T. G. O’connor. 2000. Elephant utilization of *Colophospermum mopane*: possible benefits of hedging. African Journal of Ecology 38: 352–359. doi:10.1046/j.1365-2028.2000.00258.x.

Sollmann, R., M. M. Furtado, B. Gardner, H. Hofer, A. T. A. Jácomo, N. M. TÔrres, and L. Silveira. 2011/3. Improving density estimates for elusive carnivores: Accounting for sex-specific detection and movements using spatial capture–recapture models for jaguars in central Brazil. Biological conservation 144: 1017–1024.

Sunquist, M., and F. Sunquist. 2002. Wild Cats of the World. University of Chicago Press.

Taylor, P. J. 2013. *Otomys irroratus* Southern African vlei rat. in Mammals of Africa: Rodents, Hares and Rabbits, ed. D. C. D. Happold, 583–585. London: Bloomsbury. doi:10.5040/9781472926937.0355.

Thiel, C. 2011. Ecology and population status of the Serval Leptailurus serval (Schreber, 1776) in Zambia. PhD Thesis, University of Bonn.

Thiel, C. 2015. Leptailurus serval. The IUCN Red List of Threatened Species 2015: e.T11638A50654625. doi:10.2305/IUCN.UK.2015-2.RLTS.T11638A50654625.en.

Tobler, M. W., and G. V. N. Powell. 2013. Estimating jaguar densities with camera traps: Problems with current designs and recommendations for future studies. Biological Conservation 159: 109–118. doi:10.1016/j.biocon.2012.12.009.

United Nations Environment Programme. 2012. Global environmental outlook 5: environment for the future we want. United Nations Environment Programme, Valletta.

Wegge, P., C. P. Pokheral, and S. R. Jnawali. 2004. Effects of trapping effort and trap shyness on estimates of tiger abundance from camera trap studies. Animal Conservation 7: 251-256. doi: 10.1017/S1367943004001441.

de Wet, F. 2016. Veld condition assessment and management within Sasol grasslands. EnviroPulse, Hilton.

Wiesel, I. 2010. Killing of Cape fur seal (*Arctocephalus pusillus pusillu*) pups by brown hyenas (*Parahyaena brunnea*) at mainland breeding colonies along the coastal Namib Desert. Acta Ethologica 13: 93–100. doi:10.1007/s10211-010-0078-1.

Williams, N., M. W. Schwartz, and P. A. Vesk. 2009. A conceptual framework for predicting the effects of urban environments on floras. Journal of Ecology 97: 4–9. doi: 10.1111/j.1365-2745.2008.01460.x.

Williams, S. T., N. Maree, P. Taylor, S. R. Belmain, M. Keith, and L. H. Swanepoel. 2018. Predation by small mammalian carnivores in rural agro-ecosystems: An undervalued ecosystem service? Ecosystem Services. doi:10.1016/j.ecoser.2017.12.006.

Zarfl, C., A. E. Lumsdon, J. Berlekamp, L. Tydecks, and K. Tockner. 2014. A global boom in hydropower dam construction. Aquatic Sciences 77: 161–170. doi:10.1007/s00027-014-0377-0.

